# Psychotic symptoms in 16p11.2 copy number variant carriers

**DOI:** 10.1101/621250

**Authors:** Amandeep Jutla, J. Blake Turner, LeeAnne Green Snyder, Wendy K. Chung, Jeremy Veenstra-VanderWeele

## Abstract

16p11.2 copy number variation (CNV) is implicated in neurodevelopmental disorders, with the duplication and deletion associated with autism spectrum disorder (ASD) and the duplication associated with schizophrenia (SCZ). The 16p11.2 CNV may therefore provide insight into the relationship between ASD and SCZ, distinct disorders that co-occur at an elevated rate and are difficult to distinguish from each other and from common co-occurring diagnoses such as obsessive compulsive disorder (OCD), itself a potential risk factor for SCZ. As psychotic symptoms are core to SCZ but distinct from ASD, we sought to examine their predictors in a population (n = 546) of 16p11.2 CNV carriers and their noncarrier siblings recruited by the Simons Variation in Individuals Project. We hypothesized that psychotic symptoms would be most common in duplication carriers followed by deletion carriers and noncarriers, that an ASD diagnosis would predict psychotic symptoms among CNV carriers, and that OCD symptoms would predict psychotic symptoms among all participants. Using data collected across multiple measures, we identified 19 participants with psychotic symptoms. Logistic regression models adjusting for biological sex, age, and IQ found that 16p11.2 duplication and ASD diagnosis predicted psychotic symptom presence. Our findings suggest that the association between 16p11.2 duplication and psychotic symptoms is independent of ASD diagnosis and that ASD diagnosis and psychotic symptoms may be associated in 16p11.2 CNV carriers.

**Lay Summary:** Either deletion or duplication at chromosome 16p11.2 raises the risk of autism spectrum disorder, and duplication, but not deletion, has been reported in schizophrenia. In a sample of 16p11.2 deletion and duplication carriers, we found that having the duplication or having an autism diagnosis may increase the risk of psychosis, a key feature of schizophrenia.

## 1 Introduction

Copy number variation (CNV) is a type of structural genetic variation involving the deletion or duplication of a DNA segment. CNVs are common, often benign, and represent an important mechanism by which humans maintain genetic diversity (Zarrei, MacDonald, Merico, & Scherer, 2015). However, certain specific CNVs are associated with pathology, including neuropsychiatric conditions (Cook Jr & Scherer, 2008). One such CNV is the BP4-BP5 16p11.2 copy number variant (CNV), which involves approximately 600 kilobases and 29 genes (Simons VIP Consortium, 2012). Though rare in the general population, it is overrepresented in those with developmental delay or psychiatric illness. In particular, both the 16p11.2 deletion and duplication are associated with autism spectrum disorder (ASD) (Weiss et al., 2008), and 16p11.2 duplication is associated with schizophrenia (SCZ) (Kushima et al., 2018; Marshall et al., 2017; McCarthy et al., 2009). ASD prevalence is thought to be similar in both groups, with SCZ symptoms more common in duplication than deletion carriers (Niarchou et al., 2019). The 16p11.2 CNV may provide insight into the complex relationship between symptoms of ASD and symptoms of SCZ, which, while considered distinct psychiatric disorders, converge at the levels of diagnosis, neurodevelopment and epidemiology.

At a diagnostic level, ASD and SCZ share features. In ASD, impaired social-emotional reciprocity is a requirement for the diagnosis (Lord, Elsabbagh, Baird, & Veenstra-VanderWeele, 2018). In SCZ, psychosis is the disorder’s hallmark, and can be defined as a gross impairment in the ability to distinguish between inner experience and external reality (Lieberman & First, 2018). “Psychotic symptoms,” which include delusional beliefs and perceptual disturbances, reflect this impairment, and are quite distinct from ASD. However, another core SCZ feature, the so-called “negative symptoms,” include diminished emotional expression and asociality, and share many features with ASD’s social impairment (Hommer & Swedo, 2015).

The nosology of ASD and SCZ in fact has a long and complicated history (J. Rapoport, Chavez, Greenstein, Addington, & Gogtay, 2009; Wolff, 2004).

In 1910, Bleuler originally coined the term “autism” to describe the “withdrawal of the patient to his fantasies” in schizophrenia (Kuhn & Cahn, 2004). Subsequently, Kanner (1943) used the same word to describe the “extreme aloneness from the very beginning of life” in a group of children who, he surmised, had a syndrome that was separate from but related to schizophrenia. For decades thereafter, Kanner’s syndrome, variously called “infantile autism” and “infantile psychosis,” was considered one of “the childhood schizophrenias.” By the early 1970s, however, mounting evidence suggested that autism and schizophrenia were distinct disorders (Kolvin, 1971; Rutter, 1972 Oct-Dec). In 1980, this distinction was formally codified (American Psychiatric Association, 1980).

It has, however, long been recognized that subtle symptoms, such as delay and abnormality in language, often precede the emergence of frank psychotic behavior (Courvoisie, Labellarte, & Riddle, 2001; Millan et al., 2016), and SCZ increasingly has been considered a disorder of abnormal neurodevelopment (Insel, 2010; Owen, O’Donovan, Thapar, & Craddock, 2011; J. L. Rapoport, Giedd, & Gogtay, 2012). A recent meta-analysis showed that ASD and SCZ co-occur more frequently than chance would suggest, with SCZ over three times as common in individuals with ASD as in controls (Zheng, Zheng, & Zou, 2018).

If those with ASD are at elevated risk of SCZ, then recognizing psychotic symptoms in this population is of particular importance. Unlike ASD, which tends to be stable into adulthood (Lord et al., 2018), SCZ is characterized by a progressive deterioration in functioning that early detection and treatment may mitigate (Lieberman & First, 2018).

Yet the communication impairment and repetitive speech or behavior associated with ASD can make assessment and differentiation of delusional beliefs and perceptual disturbances difficult. Further, repetitive behaviors in ASD are sometimes difficult to distinguish from symptoms of obsessive compulsive disorder (OCD), which is itself a common co-occurring diagnosis that shares genetic liability with SCZ and, by extension, ASD (Consortium et al., 2018). Although OCD symptoms and characteristic repetitive behaviors in ASD are thought to be phenomeno-logically distinct (Guo et al., 2017; Jiujias, Kelley, & Hall, 2017), the boundary between them is not always clear. Obsessive compulsive symptoms may also be important in the context of recognizing psychosis. Obsessive compulsive symptoms are present in about 30% of people with SCZ (Swets et al., 2014), and recent evidence has suggested that they may represent a SCZ risk factor (Barzilay et al., 2018; Meier et al., 2014; Van Dael et al., 2011).

We sought to examine predictors of psychotic symptoms in 16p11.2 CNV carriers. By doing so, we hoped to yield insights relevant to psychosis in the broader ASD population, improving the understanding of ASD, SCZ, and the relationship these disorders have with each other and with OCD. We hypothesized that: 1) psychotic symptoms are most common in 16p11.2 duplication carriers followed by 16p11.2 deletion carriers and noncarriers, 2) the presence of an ASD diagnosis predicts an increased risk of having psychotic symptoms among CNV carriers, and 3) OCD symptoms will predict psychotic symptoms among both CNV carriers and noncarriers.

## 2 Method

### 2.1 Study Sample

Probands with the 16p11.2 CNV were identified by routine clinical testing and were recruited by the Simons Variation in Individuals Project (VIP) (Simons VIP Consortium, 2012), a large study of specific recurrent genetic variants that contribute to the risk of ASD and other neurodevelopmental disorders. Probands were recruited from across the United States and Canada. Recruitment strategies included targeted online advertising via Google and Facebook; links from patient advocacy websites; direct mailings to clinicians (such as genetic counselors, child neurologists and developmental pediatricians); and collaborations with cytogenetics laboratories. Once a proband was confirmed to have the CNV, their biological relatives had cascade genetic testing to identify additional carriers. Carriers were defined as participants with the canonical 600kb BP4-BP5 16p11.2 duplication or deletion (chromosome 16 position 29,652,999-30,199,351 in hg19). Individuals with any additional mutations known to be associated with neurodevelopmental abnormalities (including chromosomal disorders such as fragile X, other known pathogenic CNVs such as 15q11.2, or monogenic disorders such as tuberous sclerosis) were excluded. This method produced the complete Simons VIP cohort of 658 participants: 127 16p11.2 duplication (54 initially identified probands, 73 identified through cascade testing), 137 16p11.2 deletion (115 initially identified probands, 22 identified through cascade testing), and 394 noncarrier relatives. Our study included all cohort members who were evaluated for ASD and completed an IQ assessment. 546 participants met these criteria: 109 with 16p11.2 duplication (52 initially identified probands, 57 identified through cascade testing), 131 with 16p11.2 deletion (111 initially identified probands, 20 identified through cascade testing), and 306 noncarriers.

Within the study sample, we compared several baseline characteristics of 16p11.2 duplication, 16p11.2 deletion, and noncarrier participants. Mean age and IQ were compared using analysis of variance (ANOVA), with Tukey’s procedure used for post-hoc pairwise comparisons. Biological sex, ASD diagnosis, and OCD symptoms were compared using *χ*^2^, with Bonferroni-adjusted *χ*^2^ for post-hoc comparisons (**Table 1**).

**Table 1:**
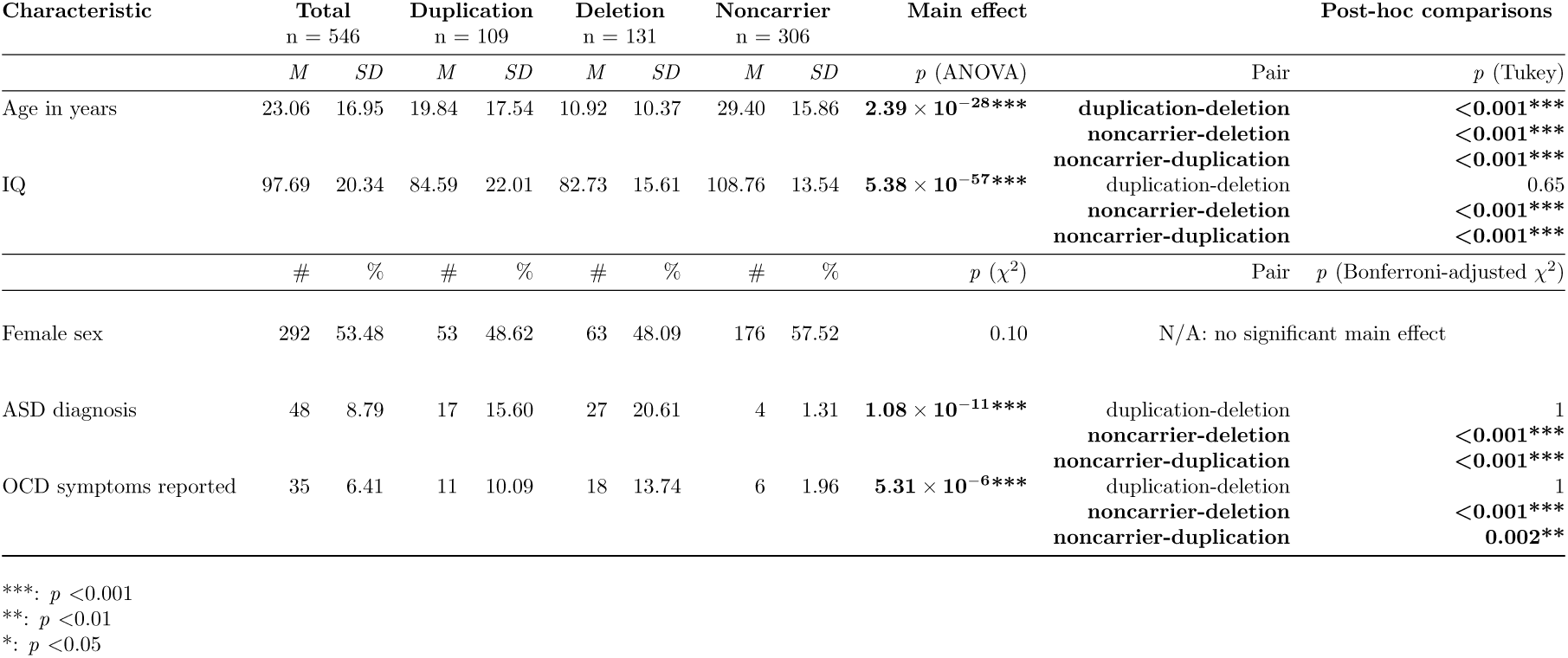
Sample characteristics

| Characteristic | Total<br>n = 546 |  | Duplication<br>n = 109 |  | Deletion<br>n = 131 |  | Noncarrier<br>n = 306 |  | Main effect | Post-hoc comparisons |  |
| --- | --- | --- | --- | --- | --- | --- | --- | --- | --- | --- | --- |
|  | *M* | *SD* | *M* | *SD* | *M* | *SD* | *M* | *SD* | *p* (ANOVA) | Pair | *p* (Tukey) |
| Age in years | 23.06 | 16.95 | 19.84 | 17.54 | 10.92 | 10.37 | 29.40 | 15.86 | **2.39 × 10<sup>-28\*\*\*</sup>** | duplication-deletion<br>noncarrier-deletion<br>noncarrier-duplication | <0.001***<br><0.001***<br><0.001*** |
| IQ | 97.69 | 20.34 | 84.59 | 22.01 | 82.73 | 15.61 | 108.76 | 13.54 | **5.38 × 10<sup>-57\*\*\*</sup>** | duplication-deletion<br>noncarrier-deletion<br>noncarrier-duplication | 0.65<br><0.001***<br><0.001*** |
|  | # | % | # | % | # | % | # | % | *p* (χ<sup>2</sup>) | Pair | *p* (Bonferroni-adjusted χ<sup>2</sup>) |
| Female sex | 292 | 53.48 | 53 | 48.62 | 63 | 48.09 | 176 | 57.52 | 0.10 | N/A: no significant main effect |  |
| ASD diagnosis | 48 | 8.79 | 17 | 15.60 | 27 | 20.61 | 4 | 1.31 | **1.08 × 10<sup>-11\*\*\*</sup>** | duplication-deletion<br>noncarrier-deletion<br>noncarrier-duplication | 1<br><0.001***<br><0.001*** |
| OCD symptoms reported | 35 | 6.41 | 11 | 10.09 | 18 | 13.74 | 6 | 1.96 | **5.31 × 10<sup>-6\*\*\*</sup>** | duplication-deletion<br>noncarrier-deletion<br>noncarrier-duplication | 1<br><0.001***<br>0.002** |

\*\*\*: *p* <0.001 \*\*: *p* <0.01 \*: *p* <0.05

### 2.2 Assessment Measures

Participants traveled to one of three phenotyping sites: Baylor College of Medicine (Houston, TX), Boston Children’s Hospital (Boston, MA) or University of Washington, Seattle (Seattle, WA). Travel expenses were paid to limit financial barriers. Participants underwent a standardized assessment performed by trained clinicians that encompassed self-report, parent-report, interview, and observation measures, with the measures a particular participant received varying based on age and carrier status (**Table 2**).

**Table 2:** Phenotypic assessment measures

| Domain | Measure | Age | Type | Total<br>n = 546 | Duplication<br>n = 109 | Deletion<br>n = 131 | Noncarrier<br>n = 306 |
| --- | --- | --- | --- | --- | --- | --- | --- |
| ASD | ADOS | Youth and Adults | Clinician assessment of participant | 315 | 97 | 121 | 97 |
|  | ADI-R | Youth and Adults | Interview with parent | 116 | 33 | 74 | 9 |
|  | BAPQ | Adults | Questionnaire (participant) | 252 | 36 | 13 | 203 |
|  | SCQ | Youth | Questionnaire (parent) | 237 | 60 | 102 | 75 |
|  | SRS | Youth | Questionnaire (parent) | 237 | 60 | 101 | 76 |
|  | SRS-ARV | Adults | Questionnaire (individual who knows participant well) | 253 | 39 | 12 | 202 |
| IQ | Mullen | Youth | Clinician assessment of participant | 63 | 22 | 30 | 11 |
|  | DAS-II Early Years (Lower) | Youth | Clinician assessment of participant | 28 | 8 | 12 | 8 |
|  | DAS-II Early Years (Upper) | Youth | Clinician assessment of participant | 60 | 13 | 24 | 23 |
|  | DAS-II School Age | Youth | Clinician assessment of participant | 151 | 35 | 65 | 51 |
|  | WASI | Adults | Clinician assessment of participant | 271 | 42 | 14 | 215 |
| Psychiatric symptoms | CBCL | Youth | Questionnaire (parent) | 194 | 47 | 85 | 62 |
|  | ABCL | Adults | Questionnaire (individual who knows participant well) | 88 | 37 | 12 | 39 |
|  | SCL-90-R | Adults | Questionnaire (participant) | 271 | 43 | 14 | 214 |
|  | DISC | Youth | Interview with parent | 178 | 42 | 81 | 55 |
|  | M-SOPS | Youth and Adults | Clinician assessment of subject | 26 | 15 | 8 | 3 |

ASD diagnoses were made based on clinical judgment informed by the results of clinician-administered and self- or caregiver-report measures. The Autism Diagnostic Observation Scale, Second Edition (ADOS-2) (Lord et al., 2012), a clinician-administered observational measure, was administered to all participants except noncarrier parents of carrier children or participants in whom the measure’s use was not feasible due to limitations of cognition or mobility. An ADOS-2 assessment involves the administration of one of four modules designed for different levels of verbal ability and, in the case of Module 4, age. Raw scores are produced for core domains of social affect (SA) and restricted/repetitive behaviors (RRB), as well as a combined “total” raw score for overall ASD symptomatology. These raw scores can be converted into scaled “Calibrated Severity Scores” (CSS) that range from 1 to 10 and represent a standardized quantification of ASD symptom severity (Gotham, Pickles, & Lord, 2009; Hus, Gotham, & Lord, 2014; Hus & Lord, 2014). The Autism Diagnostic Interview-Revised (ADI-R) (Rutter, Le Couteur, & Lord, 2003), an interview with the participant’s parent or caregiver, was administered to all participants in whom ASD was suspected. Self-or caregiver-report measures were also used to inform the clinical ASD diagnosis, including the Broad Autism Phenotype Questionnaire (BAPQ) (Hurley, Losh, Parlier, Reznick, & Piven, 2007), Social Communication Questionnaire (SCQ) (Rutter, Bailey, & Lord, 2003) and Social Responsiveness Scale (SRS)/Social Responsiveness Scale-Adult Research Version (SRS-ARV) (Constantino, 2005; Constantino & Todd, 2005).

IQ was measured with the Differential Ability Scales, Second Edition (DAS-II) (Elliot, 2007) and Mullen Scales of Early Learning (MSEL) (Shank, 2011) in children and the Wechsler Abbreviated Scale of Intelligence (WASI) (Wechsler, 1999) in adults. Adaptive skills were assessed using the Vineland Adaptive Behavior Scales II (Sparrow, Cicchetti, & Balla, 2005). Psychiatric symptoms were assessed using the school-age Child Behavior Checklist (CBCL), Adult Behavior Checklist (ABCL), Symptom Checklist-90-Revised (SCL-90-R), DISC (Diagnostic Interview Schedule for Children), and M-SOPS (Modified Scale of Prodromal Symptoms). The CBCL is part of the Achenbach System of Empirically Based Assessment (ASEBA), and consists of 113 questions about mental health with eight underlying factors (Achenbach & Rescorla, 2001). It is normed for six to eighteen-year-olds and completed by a parent or caregiver. The ABCL is an analogous ASEBA scale for adults, normed for ages eighteen to 59 and completed by an adult who knows the participant well (Achenbach & Rescorla, 2003). The SCL-90-R is a 90-item Likert-type self-report measure of psychiatric symptoms in adults, with nine underlying factors (Derogatis, 1994). The DISC is a structured diagnostic interview designed to assess for symptoms of DSM-IV psychiatric disorders in children and adolescents (Shaffer, Fisher, Lucas, Dulcan, & Schwab-Stone, 2000). The M-SOPS is a nineteen-item clinician-rated instrument that measures symptoms of psychosis (McGlashan, Miller, Woods, Hoffman, & Davidson, 2001).

### 2.3 Analytic Approach

#### 2.3.1 Development of a Psychotic Symptom Index

A psychosis-specific measure, the M-SOPS, was only administered to 26 participants. We therefore derived a composite index of psychotic symptoms by combining M-SOPS responses with data collected from the CBCL/ABCL, SCL-90-R, and DISC, which all include questions assessing for psychotic symptoms **(Table S1)**. 463 (84.80%) participants received at least one of these four measures, and 276 (50.55%) received two or more.

For each measure, we derived a binary variable indicating a screen-positive or negative for presence/absence of psychotic symptoms based on predefined criteria. Then, for each pairwise combination of measures, we examined the extent to which positive screens co-occurred and performed Fisher’s exact test to assess the strength of their relationship.

If a subject screened positive by at least two different measures, we considered the composite index to be positive, reflecting the likely presence of psychotic symptoms. To interrogate the robustness of this indicator, we created and compared four versions of the composite index. Version one, which we created first, was the least stringent. Version two used an age cutoff, version three used a stricter CBCL/ABCL threshold, and version four incorporated both.

Positive screens by each measure comprising the index were operationalized as follows:

##### CBCL/ABCL

The CBCL/ABCL “Thought Problems” factor includes several psychosis-related items. As item-level CBCL/ABCL data were not available, for version one of the index we selected a Thought Problems T-score threshold of ≥ 60 to identify scores at least one standard deviation above the mean, and considered these positive. As the CBCL Thought Problems T-Score can be elevated in nonpsychotic youth with ASD (Biederman et al., 2010; Duarte, Bordin, de Oliveira, & Bird, 2003; Hoffmann, Weber, König, Becker, & Kamp-Becker, 2016; Mazefsky, Anderson, Conner, & Minshew, 2011; Ooi, Rescorla, Ang, Woo, & Fung, 2011), versions three and four of the index raised the threshold to ≥ 70 (i.e., two rather than one standard deviations above the mean).

##### SCL-90-R

We selected four items reflecting specific psychotic symptoms distinct from ASD from the SCL-90-R “psychoticism” factor: “the idea that someone else can control your thoughts,” “hearing voices that other people do not hear,” “other people being aware of your private thoughts,” and “having thoughts that are not your own.” We considered a response of at least “a little bit” to any of these items to be a positive screen.

##### DISC

For each DSM-IV diagnosis assessed by the DISC interview, data were available regarding the number of symptoms endorsed but not which were endorsed specifically. We considered endorsement of at least one schizophrenia symptom within the past year to represent a positive screen.

##### M-SOPS

Five M-SOPS items assess symptoms of psychosis: “unusual thought content or delusional ideas,” “suspiciousness or persecutory ideas,” “grandiosity,” “perceptual abnormalities or hallucinations,” and “disorganized communication.” The presence of at least one of these symptoms (with the exception of “disorganized communication,” which we did not consider given its non-specificity) represented a positive screen.

Versions one and three of the index did not incorporate an age cutoff. However, since true psychosis in young children is rare, with childhood-onset schizophrenia typically not presenting before age seven (Baribeau & Anagnostou, 2013), versions two and four required that a participant be at least seven years old to be positively identified with psychotic symptoms.

#### 2.3.2 Primary Analysis

As index version four was the most stringent, incorporating both the raised CBCL threshold and the age cutoff, we used it to identify participants likely to have psychotic symptoms. We then examined predictors of the presence of psychotic symptoms by conducting a series of logistic regressions. All models used generalized estimating equations (GEEs) to control for intra-family correlations (Hanley, Negassa, deB Edwardes, & Forrester, 2003).

Our predictor variables of interest, which we selected *a priori*, were CNV carrier status, age, IQ, clinical ASD diagnosis, OCD symptoms (as measured by endorsement of at least one OCD symptom in the past year during the DISC interview) and biological sex. Prior to conducting any analyses, we ruled out multicollinearity by inspecting the correlation matrix between scaled versions of all variables.

Our primary analysis included four regression models. The first was estimated for the entire sample, and included all predictors of interest. The second, third and fourth models were estimated for subgroups of the sample defined by carrier status (i.e., 16p11.2 deletion carriers, 16p11.2 duplication carriers, and noncarriers), and each included all predictors of interest except carrier status. All analyses used unscaled variables for ease of interpretability.

#### 2.3.3 Exploratory Regression Analyses

To determine whether ASD severity could predict the presence of psychotic symptoms, we estimated exploratory regression models that substituted the categorical ASD diagnosis predictor with continuous ADOS CSS values.

Total CSS values for participants who received ADOS Modules 1, 2 or 3 were available to us as part of the Simons VIP dataset. For those who received ADOS Module 4, we derived total CSS values from item-level data (Hus & Lord, 2014). For all ADOS modules, we derived SA and RRB domain CSS values from item-level data where available (Hus et al., 2014). In exploratory models for which total CSS was a predictor, we excluded participants who did not receive the ADOS, yielding a reduced sample (total n = 315, with 97 duplication carriers, 121 deletion carriers, and 97 noncarriers). In exploratory models for which domain CSS values were predictors, we excluded participants who lacked item-level data, and whose domain scores therefore could not be derived. This reduced the sample further (total n = 249, with 68 duplication carriers, 97 deletion carriers, and 82 noncarriers).

#### 2.3.4 Software and Data

We conducted all analyses in R 3.5.1 (R Core Team, 2018), using functions from dplyr 0.7.8 (Wickham, François, Henry, & Müller, 2018), magrittr 1.5 (Wickham & Bache, 2014), and purrr 0.2.5 (Henry & Wickham, 2019), as well as *chisq.post.hoc* from fifer 1.1 (Fife, 2014, March 28/ 2019), *rescale* from arm 1.10-1 (Gelman et al., 2018), *geeglm* from geepack 1.2-1 (Hojsgaard, Halekoh, & Yan, 2016), and *tidy* from broom 0.5.0 (Robinson et al., 2018). Analysis scripts are available from the authors at https://github.com/amandeepjutla/2019-16p11-psychosis. The Simons VIP 16p11.2 v10.0 dataset used for this study can be requested through the Simons Foundation Autism Research Initiative (SFARI, RRID:SC 004261) online portal, SFARI Base, at https://base.sfari.org.

## 3 Results

### 3.1 Sample Characteristics

The sample represented a broad range of ages (*M* = 23.06, *SD* = 16.95 years), with significant variation in age among 16p11.2 duplication, 16p11.2 deletion, and noncarriers, *F* (2, 543) = 71.67, *p <* 2.39×10^−28^, and post-hoc comparisons showed significant differences for duplication-deletion, noncarrier-duplication, and noncarrier-deletion pairwise comparisons. IQ (*M* = 97.69, *SD* = 20.34) also varied significantly among the three groups, *F* (2, 543) = 166.04, *p <* 4.38 × 10^−57^, with post-hoc comparisons showing that duplication and deletion group IQ scores differed from the noncarrier group, but not from each other.

The three groups were not significantly imbalanced in terms of biological sex composition, *χ*^2^(1) = 4.57, *p* = 0.10. They differed in terms of ASD diagnosis, *χ*^2^(1) = 50.49, *p* = 1.08×10^−11^ and presence of OCD symptoms, *χ*^2^(1) = 24.29, *p* = 5.31 × 10^−6^. Post-hoc comparisons for ASD and OCD showed that, as with IQ, duplication and deletion carriers differed significantly from noncarriers but not each other.

#### 3.1.1 Participants with Psychotic Symptoms

56 of 282 participants screened positive on the CBCL or ABCL (using the ≥ 70 T-Score cutoff), 50 of 271 on SCL-90-R, 23 of 178 on DISC, and 9 of 26 on M-SOPS **(Table 3)**. We observed some degree of overlap for all possible pairwise combinations of these measures except SCL-90 × DISC, which was expected because SCL-90 was given only to adults and DISC only to children. Tests of relationship strength between pairs (**Table 4**) identified a statistically significant association between CBCL/ABCL × DISC (OR 7.71, 95% CI 2.16 - 42.21, *p* = 2.29 × 10^−4^).

**Table 3:** Index measures by carrier status

| Measure | Total<br>n = 546 |  |  | Duplication<br>n = 109 |  |  | Deletion<br>n = 131 |  |  | Noncarrier<br>n = 306 |  |  |
| --- | --- | --- | --- | --- | --- | --- | --- | --- | --- | --- | --- | --- |
|  | # Received | # Positive | % Positive | # Received | # Positive | % Positive | # Received | # Positive | % Positive | # Received | # Positive | % Positive |
| CBCL/ABCL | 282 | 56 | 19.86 | 84 | 27 | 32.14 | 97 | 21 | 21.65 | 101 | 8 | 7.92 |
| SCL-90-R | 271 | 50 | 18.45 | 43 | 19 | 44.19 | 14 | 7 | 50 | 214 | 24 | 11.21 |
| DISC | 178 | 23 | 12.92 | 42 | 7 | 16.67 | 81 | 8 | 9.88 | 55 | 8 | 14.55 |
| SOPS | 26 | 9 | 5.06 | 15 | 5 | 11.9 | 8 | 3 | 3.7 | 3 | 1 | 1.82 |

**Table 4:**
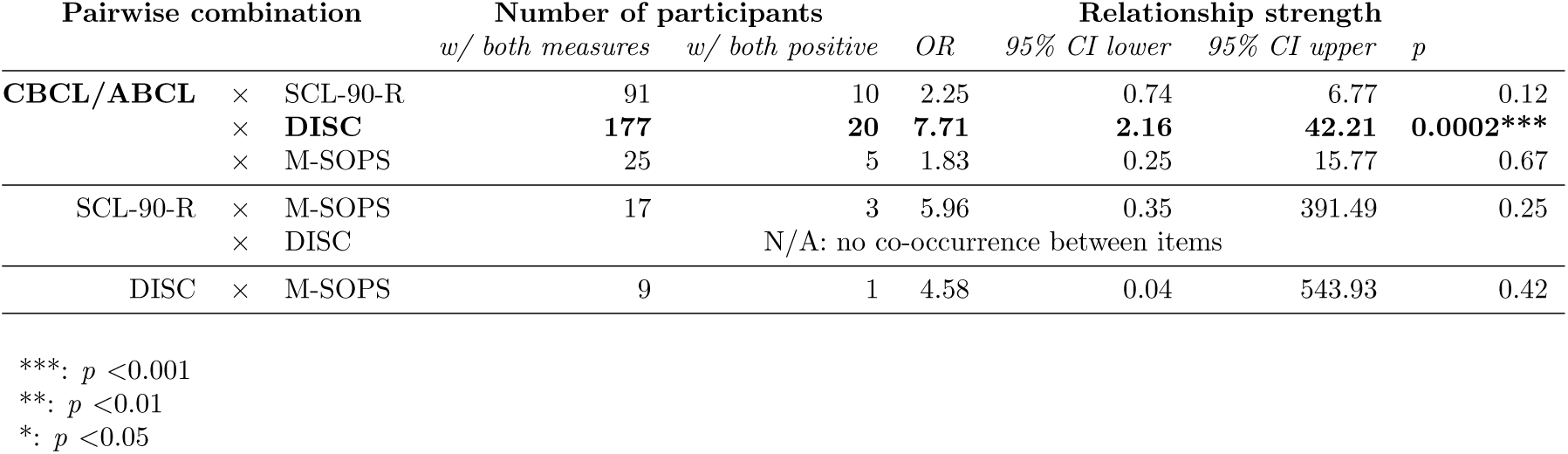
Pairwise combinations between index measures

| Pairwise combination |  |  | Number of participants |  | Relationship strength |  |  |  |
| --- | --- | --- | --- | --- | --- | --- | --- | --- |
|  |  |  | *w/ both measures* | *w/ both positive* | *OR* | *95% CI lower* | *95% CI upper* | *p* |
| CBCL/ABCL | × | SCL-90-R | 91 | 10 | 2.25 | 0.74 | 6.77 | 0.12 |
|  | × | DISC | **177** | **20** | **7.71** | **2.16** | **42.21** | **0.0002\*\*\*** |
|  | × | M-SOPS | 25 | 5 | 1.83 | 0.25 | 15.77 | 0.67 |
| SCL-90-R | × | M-SOPS | 17 | 3 | 5.96 | 0.35 | 391.49 | 0.25 |
|  | × | DISC | N/A: no co-occurrence between items |  |  |  |  |  |
| DISC | × | M-SOPS | 9 | 1 | 4.58 | 0.04 | 543.93 | 0.42 |

\*\*\*: $p < 0.001$
\*\*: $p < 0.01$
\*: $p < 0.05$

Using the most stringent version of the composite index (version four), nineteen participants had likely psychotic symptoms. Of these, nine were female and ten were male. Twelve had 16p11.2 duplication, four had 16p11.2 deletion, and three were noncarrier family members. Seven had a clinical ASD diagnosis, and three had OCD symptoms. Two, both duplication carriers, came from the same family. Most participants who met the “likely psychotic symptoms” threshold (eleven of the nineteen) did so by a combination of positive screens on the CBCL/ABCL and DISC measures. Of the remainder, three screened positive on CBCL/ABCL and SCL-90-R, three on SCL-90-R and M-SOPS, one on CBCL/ABCL and M-SOPS, and one on DISC and M-SOPS. No participant screened positive on more than two measures.

Their mean age was 18.03 years (*SD* = 10.93 years), and mean IQ was 81.95 (*SD* = 19.75).

### 3.2 Predictors of Psychotic Symptoms

The parameters of regression models estimated for the primary analysis are presented in **Table 5** (for the entire sample) and **Table 6** (for carrier status-defined subgroups).

**Table 5:** Predictors of psychotic symptoms in entire sample

| Predictor | *B* | *SE* | Wald $\chi^2$ | *OR* | *95% CI lower* | *95% CI upper* | *p* |
| --- | --- | --- | --- | --- | --- | --- | --- |
| (Intercept) | -3.98 | 1.45 | 7.57 | 0.02 | 0.00 | 0.32 | 0.01 |
| **Duplication** | **2.01** | **0.73** | **7.52** | **7.44** | **1.77** | **31.18** | **0.006\*\*** |
| Deletion | 0.51 | 0.89 | 0.32 | 1.66 | 0.29 | 9.55 | 0.57 |
| Age in years | 0.01 | 0.01 | 0.37 | 1.01 | 0.98 | 1.03 | 0.55 |
| IQ | -0.01 | 0.01 | 0.53 | 0.99 | 0.97 | 1.02 | 0.47 |
| **ASD diagnosis** | **1.44** | **0.6** | **5.81** | **4.21** | **1.31** | **13.56** | **0.02\*** |
| OCD symptoms | 0.73 | 0.74 | 0.97 | 2.08 | 0.49 | 8.91 | 0.33 |
| Biological sex | 0.01 | 0.47 | 0.00 | 1.01 | 0.40 | 2.53 | 0.98 |

\*\*\*: $p < 0.001$
\*\*: $p < 0.01$
\*: $p < 0.05$

**Table 6:** Predictors of psychotic symptoms within carrier status-defined subsets

| Predictor | *B* | SE | Wald $\chi^2$ | OR | 95% CI lower | 95% CI upper | *p* |
| --- | --- | --- | --- | --- | --- | --- | --- |
| *Duplication carriers only* |  |  |  |  |  |  |  |
| (Intercept) | -1.79 | 1.52 | 1.39 | 0.17 | 0.01 | 3.26 | 0.24 |
| Age in years | 0.02 | 0.02 | 1.29 | 1.02 | 0.99 | 1.05 | 0.26 |
| IQ | -0.01 | 0.02 | 0.41 | 0.99 | 0.96 | 1.02 | 0.52 |
| ASD diagnosis | 1.49 | 0.81 | 3.4 | 4.46 | 0.91 | 21.81 | 0.07 |
| OCD symptoms | N/A : no duplication carriers positive for psychotic symptoms had OCD symptoms |  |  |  |  |  |  |
| Biological sex | -0.29 | 0.68 | 0.18 | 0.75 | 0.2 | 2.85 | 0.67 |
| *Deletion carriers only* |  |  |  |  |  |  |  |
| (Intercept) | -6.52 | 3.62 | 3.25 | 0.00 | 0.00 | 1.76 | 0.07 |
| Age in years | 0.00 | 0.03 | 0.02 | 1.00 | 0.95 | 1.06 | 0.90 |
| IQ | 0.02 | 0.03 | 0.46 | 1.02 | 0.96 | 1.08 | 0.50 |
| ASD diagnosis | 1.41 | 1.17 | 1.45 | 4.10 | 0.41 | 40.63 | 0.23 |
| OCD symptoms | 1.94 | 1.17 | 2.76 | 6.99 | 0.70 | 69.31 | 0.10 |
| Biological sex | 0.47 | 1.24 | 0.14 | 1.60 | 0.14 | 18.05 | 0.70 |
| *Noncarriers only* |  |  |  |  |  |  |  |
| (Intercept) | -0.71 | 2.91 | 0.06 | 0.49 | 0 | 146.86 | 0.81 |
| **Age in years** | **-0.08** | **0.03** | **5.58** | **0.93** | **0.87** | **0.99** | **0.02** |
| IQ | -0.03 | 0.03 | 0.85 | 0.97 | 0.91 | 1.03 | 0.36 |
| ASD diagnosis | N/A: no noncarriers positive for psychotic symptoms had ASD |  |  |  |  |  |  |
| OCD symptoms | 2.09 | 1.46 | 2.05 | 8.12 | 0.46 | 142.96 | 0.15 |
| Biological sex | 0.56 | 1.69 | 0.11 | 1.75 | 0.06 | 47.71 | 0.74 |

\*\*\*: $p < 0.001$
\*\*: $p < 0.01$
\*: $p < 0.05$

#### 3.2.1 Hypothesis 1: CNV Carrier Status as Predictor

Hypothesis 1, that psychotic symptoms would be most common in 16p11.2 duplication carriers followed by 16p11.2 deletion carriers and noncarriers was partially supported by our finding that, in the model estimated for the entire sample, 16p11.2 duplication carrier status predicted psychotic symptom presence (OR 7.44, 95% CI 1.77 - 31.18, *p* = 0.006). Neither deletion carrier status nor noncarrier status was a significant predictor.

#### 3.2.2 Hypothesis 2: ASD Defined by Clinical Diagnosis as Predictor

Hypothesis 2, that ASD diagnosis would predict presence of psychotic symptoms among 16p11.2 CNV carriers, was partially supported by our finding that categorical ASD diagnosis predicted psychotic symptom presence in the entire sample (OR 4.21, 95% CI 1.31 - 13.56, *p* = 0.02). An insufficient number of noncarriers had an ASD diagnosis, or co-occurring psychotic symptoms, to interpret findings against other subgroups. ASD diagnosis did not reach statistical significance as a predictor among either CNV carrier-defined subgroup alone.

#### 3.2.3 Hypothesis 3: OCD Symptoms as Predictor

Hypothesis 3, that OCD symptoms would predict the presence of psychotic symptoms among both carriers and noncarriers, was not significantly supported by our findings.

#### 3.2.4 IQ, Biological Sex and Age as Predictors

IQ and biological sex were not significant predictors of the presence of psychotic symptoms in the entire sample or any of its subgroups. Age reached statistical significance as a negative predictor among noncarriers (OR 0.93 for every year increase in age, 95% CI 0.87 - 0.99, *p* = 0.02). This is consistent with evidence that, in the neurotypical population, hallucinations are more common in children than adults (Maijer, Begemann, Palmen, Leucht, & Sommer, 2018). However, as only three noncarriers had psychotic symptoms, this finding is likely to be artifactual.

#### 3.2.5 Exploration of ASD Severity as Predictor

The parameters of exploratory models that substituted categorical ASD diagnosis with continuous ADOS Calibrated Severity Scores (CSS) are presented in **Table S2** (for total CSS) and **Table S3** (for domain CSS).

Total CSS trended toward significance as a predictor of psychotic symptoms among all participants who received the ADOS (OR 1.21 for every one point increase in CSS, 95% CI 0.99-1.47, *p* = 0.06). We did not find that domain CSS for RRB or SA were significant predictors.

#### 3.2.6 Robustness of Findings

Less stringently-defined versions of the composite psychotic symptom index produced results similar to the version four results reported above. Duplication status and ASD diagnosis consistently predicted psychotic symptoms.

Version one, which had a CBCL/ABCL T-Score threshold of ≥ 60 and no age cutoff, identified thirty-five participants as having likely psychotic symptoms. Using this group, duplication status, ASD diagnosis, and OCD symptoms were significant predictors of psychotic symptoms in the entire sample (duplication: OR 5.13, 95% CI 1.70 - 15.49, *p <* 0.001; ASD diagnosis: OR 2.83, 95% CI 1.08 - 7.40, *p* = 0.03; OCD symptoms: OR 3.32, 95% CI 1.14 - 9.70, *p* = 0.03). OCD symptoms were also a significant predictor among deletion carriers alone (OR 7.22, 95% CI 1.30 - 40.09, *p* = 0.02).

Version two, which added the requirement that a participant to be at least seven years old to be identified with psychotic symptoms, reduced the number identified from thirty-five to thirty. Here, duplication status and ASD diagnosis, but not OCD, were significant predictors of psychotic symptoms in the entire sample (duplication: OR 6.29, 95% CI 1.86 - 21.25, *p <* 0.01, ASD: OR 2.80, 95% CI 1.02 - 7.70, *p* = 0.046).

Version three, which had no age cutoff but raised the CBCL/ABCL threshold, reduced participants identified as likely having psychotic symptoms from thirty-five to twenty-one. Duplication status and ASD continued to predict psychotic symptoms in the entire sample (duplication: OR 6.64, 95% CI 1.81 - 24.39, *p <* 0.01; ASD: OR 4.13, 95% CI 1.27 - 13.37, *p* = 0.02). OCD was not statistically significant.

As the deletion carrier group was younger than the duplication carrier or noncarrier groups, we conducted an additional sensitivity analysis that constrained the entire sample to participants who were at least twelve years old. In this restricted sample (total n = 327, with 37 deletion carriers, 53 duplication carriers, and 237 non-carriers), we found that duplication status remained a significant predictor of psychotic symptoms (OR 39.00, 95% CI 7.34 – 208, *p* = 0.00002). ASD was not a significant predictor, and no participants in this age-constrained sample had OCD symptoms.

## 4 Discussion

Our findings indicate an association between 16p11.2 duplication status and psychotic symptoms. This aligns with previous studies that reported the 16p11.2 duplication in schizophrenia genetic samples (Giaroli, Bass, Strydom, Rantell, & McQuillin, 2014; McCarthy et al., 2009; Rees et al., 2014; Steinberg et al., 2014).

We were unable to detect a significant association between 16p11.2 deletion and psychotic symptoms. This conflicts with reports of schizophrenia diagnosis in association with 16p11.2 deletion carriers (Kushima et al., 2018; Marshall et al., 2017). However, we may have been underpowered to detect an association. Only four deletion carriers had psychotic symptoms in our sample, compared with twelve duplication carriers, which is consistent with recent evidence suggesting that psychotic symptoms may be less common in 16p11.2 deletion than duplication carriers (Niarchou et al., 2019).

Independent of the type of CNV, ASD diagnosis was also a significant predictor of psychosis risk among 16p11.2 CNV carriers in our primary analysis. Our exploratory analyses of potential relationships between ASD severity as measured by ADOS Calibrated Severity Scores and psychosis risk did not yield significant results. However, many participants, most of whom were noncarriers, did not receive the ADOS and had to be excluded from these models. This reduction in sample size, along with the exclusion of noncarriers, many of whom may not have had significant ASD symptoms, could have biased us against detecting an effect.

Though we did not find an association between psychotic symptoms and OCD, we did find that OCD symptoms were more common in 16p11.2 CNV carriers than noncarriers. This suggests that 16p11.2 may warrant future exploration in genetic studies of OCD, which currently are limited (Fernandez, Leckman, & Pittenger, 2018). As of now, 16p11.2 duplication has been described in, but not specifically associated with, OCD (McGrath et al., 2014).

This study has important strengths, primarily pertaining to the unique Simons VIP sample. The specific focus on a rare genetic variant allowed us to minimize underlying genetic heterogeneity in exploring the relationship between ASD and risk of psychotic symptoms. Further, we tested convergent validity across multiple measures within our psychotic symptom index. We also were able to verify the stability of our results using alternate versions of the composite psychotic symptom index with different levels of stringency.

This study also has important limitations. Our use of the VIP cohort, despite its advantages, necessarily restricted the conclusions we could draw. Although Simons VIP sought to mitigate ascertainment bias by conducting cascade testing, a large proportion of the cohort’s deletion carriers in particular were identified probands. Thus, it is unclear to what extent our findings may generalize to deletion carriers who have not come to clinical attention. As our study excluded 16p11.2 CNV carriers with additional known mutations associated with neurodevelopmental abnormalities, it is also unclear to what extent our findings might generalize to such individuals, in whom these additional genetic variants might affect their phenotype.

Our focus on a rare CNV limited our sample size, which in turn restricted the statistical power we could achieve. The ratio between the number of participants with psychotic symptoms and the number of predictors in our regression models, while in an acceptable range (van Smeden et al., 2016; Vittinghoff & McCulloch, 2007), could have introduced a potential for overfitting, particularly in subgroup analyses, though our sensitivity analyses were partially able to address this.

The deletion carriers in our study were, on average, significantly younger than other participants. Our inability to detect an association between deletion status and psychotic symptoms should therefore not be construed as evidence of no association. Psychotic symptoms could potentially develop in members of this group as they enter adolescence and young adulthood, and although we still did not detect an association when we restricted the sample to older participants, the resultant reduction in the deletion group’s sample size limited our power.

Finally, our psychotic symptom index, though carefully developed, used a combination of self-and parent-report measures with varying levels of specificity for psychosis. As the CBCL/ABCL Thought Problems factor includes behavioral symptoms other than psychosis, it is probably the least specific measure we used, followed by DISC, which incorporates DSM-IV “negative” schizophrenia symptoms that overlap with ASD. However, with the SCL-90-R and M-SOPS, we were able to use individual items with high specificity, and M-SOPS in particular was designed specifically for the detection of psychosis. We further increased specificity by requiring participants to screen positive on two different measures for us to consider them as having likely psychotic symptoms. Still, it is conceivable that there was heterogeneity in how participants met the threshold for likely psychotic symptoms and that at least some participants identified as having symptoms by the index may not have “true” clinical psychosis. Regarding this potential issue, we consider it reassuring that the majority of participants who met the threshold (15 of 19) did so by a combination of CBCL/ABCL and some other, more specific measure, either DISC (11), SCL-90-R (3) or M-SOPS (1). Our finding of a robust association between psychotic symptoms as identified by our index and 16p11.2 duplication is also consistent with existing literature.

Our work suggests several future directions for research. In subgroup analyses, we observed that ASD predicted psychotic symptoms at trend-level within the duplication group (OR: 4.46, 95% CI 0.91 – 21.81, *p* = 0.07) but not within the deletion group. This should not be over-interpreted, but may be worth exploring further, in larger samples, to determine whether it is robust. If the ASD associated with 16p11.2 duplication but not deletion is in some sense “psychosis-prone,” it may help in understanding how and why certain individuals with ASD develop SCZ while others do not.

Our “psychotic symptom index” approach should also be further tested for validity. Is it truly measuring psychotic symptoms, or are ASD-related behaviors leading to false positives? How might it perform in a population of, for example, SCZ patients without ASD? Accurate measurement will be crucial if the relationship between ASD and psychotic risk is to be delineated.

Longitudinal exploration of symptom evolution in 16p11.2 CNV carriers into adolescence and adulthood is important and is underway. This may be of particular importance for deletion carriers. Are psychotic symptoms truly less common in this group, or was this a function of its overall young age?

We hope to answer these and other questions. by conducting in-person interviews, correlating clinical metrics with neuroimaging findings, and longitudinally following the Simons VIP cohort. In doing so, we will more deeply characterize the 16p11.2 deletion and duplication phenotypes, and help generate hypotheses and insights applicable to psychotic and other symptoms in a general ASD population.

## 5 Acknowledgments

This project was financially supported by a Whitaker Scholar in Developmental Neuropsychiatry Award to AJ funded by Marilyn and James Simons Family Giving.

We would like to express our gratitude to all families participating in the Simons Variation in Individuals Project, and to the Simons Foundation Autism Research Initiative for making this project possible.

---

**Table S1:** Psychotic symptom index measures

| Measure | Item(s) |
| --- | --- |
| CBCL/ABCL | *Thought Problems T Score $\geq 60$ based on the following:*<br>Hears sound or voices that aren't there<br>Sees things that aren't there<br>Strange behavior<br>Strange ideas<br>Can't get his/her mind off certain thoughts; obsessions<br>Repeats certain acts over and over; compulsions<br>Picks nose, skin, or other parts of body (CBCL) / Picks skin or other parts of body (ABCL)<br>Plays with own sex parts too much<br>Plays with own sex parts in public<br>Stores up too many things he/she doesn't need<br>Deliberately harms self or attempts suicide<br>Nervous movements or twitching<br>Trouble sleeping<br>Talks or walks in sleep<br>Sleeps less than most kids (CBCL) / most people (ABCL) |
| SCL-90-R | *Response of at least "a little bit" to "for the past week, how much were you bothered by . . . ":*<br>The idea that someone else can control your thoughts<br>Hearing voices that other people do not hear<br>Other people being aware of your private thoughts<br>Having thoughts that are not your own |
| DISC | *At least one DSM-IV schizophrenia symptom within the past year:*<br>Delusions<br>Hallucinations<br>Disorganized speech<br>Grossly disorganized or catatonic behavior<br>Negative symptoms |
| M-SOPS | *One or more of the following symptoms is present:*<br>Unusual thought content/delusional ideas<br>Suspiciousness/persecutory ideas<br>Grandiosity<br>Perceptual abnormalities/hallucinations |

**Table S2:** ADOS Total Calibrated Severity Score as predictor of psychotic symptoms

| **Predictor** | ***B*** | **SE** | **Wald $\chi^2$** | **OR** | **95% CI lower** | **95% CI upper** | ***p*** |
| --- | --- | --- | --- | --- | --- | --- | --- |
| (Intercept) | -2.3 | 1.59 | 2.08 | 0.10 | 0.00 | 2.29 | 0.15 |
| Duplication | 0.60 | 0.66 | 0.82 | 1.82 | 0.50 | 6.70 | 0.37 |
| Deletion | -0.86 | 0.85 | 1.02 | 0.42 | 0.08 | 2.25 | 0.31 |
| Age in years | 0.02 | 0.01 | 2.25 | 1.02 | 0.99 | 1.05 | 0.13 |
| IQ | -0.02 | 0.01 | 1.68 | 0.98 | 0.96 | 1.01 | 0.19 |
| Total CSS | 0.19 | 0.10 | 3.56 | 1.21 | 0.99 | 1.47 | 0.06 |
| OCD symptoms | -0.05 | 0.87 | 0.00 | 0.95 | 0.17 | 5.22 | 0.95 |
| Biological sex | -0.13 | 0.52 | 0.06 | 0.88 | 0.32 | 2.42 | 0.80 |

\*\*\*: $p < 0.001$
\*\*: $p < 0.01$
\*: $p < 0.05$

**Table S3:**
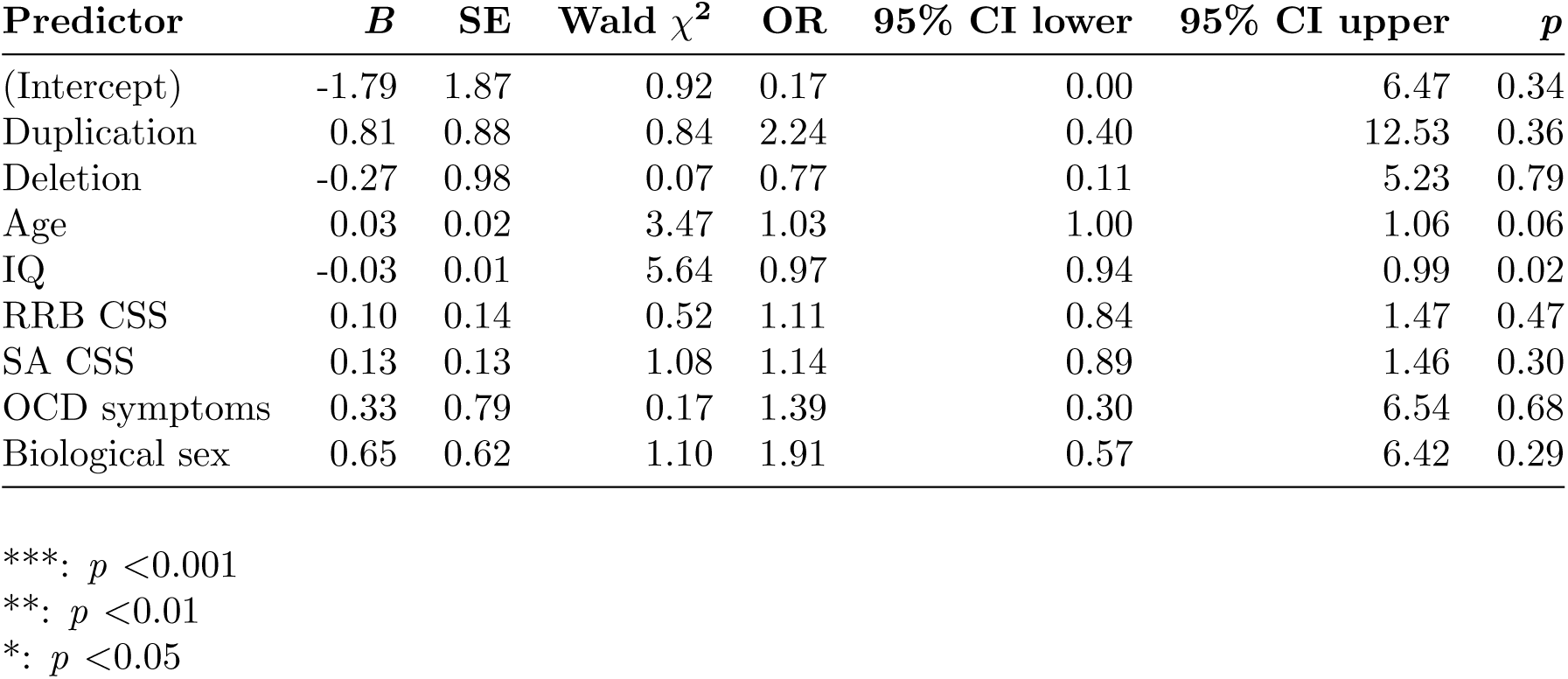
ADOS domain calibrated severity scores as predictors of psychosis

| **Predictor** | ***B*** | **SE** | **Wald $\chi^2$** | **OR** | **95% CI lower** | **95% CI upper** | ***p*** |
| --- | --- | --- | --- | --- | --- | --- | --- |
| (Intercept) | -1.79 | 1.87 | 0.92 | 0.17 | 0.00 | 6.47 | 0.34 |
| Duplication | 0.81 | 0.88 | 0.84 | 2.24 | 0.40 | 12.53 | 0.36 |
| Deletion | -0.27 | 0.98 | 0.07 | 0.77 | 0.11 | 5.23 | 0.79 |
| Age | 0.03 | 0.02 | 3.47 | 1.03 | 1.00 | 1.06 | 0.06 |
| IQ | -0.03 | 0.01 | 5.64 | 0.97 | 0.94 | 0.99 | 0.02 |
| RRB CSS | 0.10 | 0.14 | 0.52 | 1.11 | 0.84 | 1.47 | 0.47 |
| SA CSS | 0.13 | 0.13 | 1.08 | 1.14 | 0.89 | 1.46 | 0.30 |
| OCD symptoms | 0.33 | 0.79 | 0.17 | 1.39 | 0.30 | 6.54 | 0.68 |
| Biological sex | 0.65 | 0.62 | 1.10 | 1.91 | 0.57 | 6.42 | 0.29 |

\*\*\*: $p < 0.001$
\*\*: $p < 0.01$
\*: $p < 0.05$

## References

Achenbach, T. M., & Rescorla, L. (2001). Manual for the ASEBA School-Age Forms & Profiles. Burlington, VT: University of Vermont, Research Center for Children, Youth, & Families.

Achenbach, T. M., & Rescorla, L. (2003). Manual for the ASEBA Adult Forms & Profiles. Burlington, VT: University of Vermont, Research Center for Children, Youth, & Families.

American Psychiatric Association. (1980). Diagnostic and Statistical Manual of Mental Disorders (3rd). American Psychiatric Association.

Baribeau, D. A., & Anagnostou, E. (2013). A comparison of neuroimaging findings in childhood onset schizophrenia and autism spectrum disorder: A review of the literature. Frontiers in Psychiatry, 4. doi:10.3389/fpsyt.2013.00175

Barzilay, R., Patrick, A., Calkins, M. E., Moore, T. M., Wolf, D. H., Benton, T. D., … Gur, R. E. (2018, November 23). Obsessive compulsive symptomatology in community youth: Typical development or a red flag for psychopathology? Journal of the American Academy of Child & Adolescent Psychiatry, 0 (0). doi:10.1016/j.jaac.2018.06.038

Biederman, J., Petty, C. R., Fried, R., Wozniak, J., Micco, J. A., Henin, A., … Faraone, S. V. (2010, June). Child Behavior Checklist clinical scales discriminate referred youth with autism spectrum disorder: A preliminary study. Journal of Developmental & Behavioral Pediatrics, 1. doi:10.1097/DBP.0b013e3181e56ddd

Consortium, T. B., Anttila, V., Bulik-Sullivan, B., Finucane, H. K., Walters, R. K., Bras, J., … Neale, B. M. (2018, June 22). Analysis of shared heritability in common disorders of the brain. Science, 360 (6395), eaap8757. doi:10.1126/science.aap8757. pmid: 29930110

Constantino, J. N. (2005). Social Responsiveness Scale. Western Psychological Services.

Constantino, J. N., & Todd, R. D. (2005, March 15). Intergenerational transmission of subthreshold autistic traits in the general population. Biological Psychiatry, 57 (6), 655–660. doi:10.1016/j.biopsych.2004.12.014

Cook Jr, E. H., & Scherer, S. W. (2008, October 15). Copy-number variations associated with neuropsychiatric conditions. Nature. doi:10.1038/nature07458

Courvoisie, H., Labellarte, M. J., & Riddle, M. A. (2001, June). Psychosis in children: Diagnosis and treatment. Dialogues in Clinical Neuroscience, 3 (2), 79–92. pmid: 22033588

Derogatis, L. R. (1994). SCL-90-R: Symptom Checklist-90-R: Administration, scoring, and procedures manual. Pearson.

Duarte, C. S., Bordin, I. A. S., de Oliveira, A., & Bird, H. (2003, December 1). The CBCL and the identification of children with autism and related conditions in Brazil: Pilot findings. Journal of Autism and Developmental Disorders, 33 (6), 703–707. doi:10.1023/B:JADD.0000006005.31818.1c

Elliot, C. D. (2007). Differential Ability Scales-II. San Antonio, TX: Psychological Corporation.

Fernandez, T. V., Leckman, J. F., & Pittenger, C. (2018, January 1). Genetic susceptibility in obsessive-compulsive disorder. In D. H. Geschwind, H. L. Paulson, & C. Klein (Eds.), Handbook of Clinical Neurology (Vol. 148, pp. 767–781). Neurogenetics, Part II. doi:10.1016/B978-0-444-64076-5.00049-1

Fife, D. (2019, January 17). Fifer: A collection of R functions for data manipulation, data analysis, and plotting. Retrieved February 2, 2019, from https://github.com/dustinfife/fifer

Gelman, A., Su, Y.-S., Yajima, M., Hill, J., Pittau, M. G., Kerman, J., … Dorie, V. (2018, April 13). Arm: Data analysis using regression and multilevel/hierarchical models. Retrieved April 9, 2019, from https://CRAN.R-project.org/package=arm

Giaroli, G., Bass, N., Strydom, A., Rantell, K., & McQuillin, A. (2014, November 1). Does rare matter? Copy number variants at 16p11.2 and the risk of psychosis: A systematic review of literature and meta-analysis. Schizophrenia Research, 159 (2), 340–346. doi:10.1016/j.schres.2014.09.025

Gotham, K., Pickles, A., & Lord, C. (2009, May 1). Standardizing ADOS scores for a measure of severity in autism spectrum disorders. Journal of Autism and Developmental Disorders, 39 (5), 693–705. doi:10.1007/s10803-008-0674-3

Guo, W., Samuels, J. F., Wang, Y., Cao, H., Ritter, M., Nestadt, P. S., … Shugart, Y. Y. (2017, July 1). Polygenic risk score and heritability estimates reveals a genetic relationship between ASD and OCD. European Neuropsychopharmacology, 27 (7), 657–666. doi:10.1016/j.euroneuro.2017.03.011

Hanley, J. A., Negassa, A., deB Edwardes, M. D., & Forrester, J. E. (2003, February 15). Statistical analysis of correlated data using generalized estimating equations: An orientation. American Journal of Epidemiology, 157 (4), 364–375. doi:10.1093/aje/kwf215

Henry, L., & Wickham, H. (2019, March 15). Purrr: Functional programming tools. Retrieved April 9, 2019, from https://CRAN.R-project.org/package=purrr

Hoffmann, W., Weber, L., König, U., Becker, K., & Kamp-Becker, I. (2016, July 1). The role of the CBCL in the assessment of autism spectrum disorders: An evaluation of symptom profiles and screening characteristics. Research in Autism Spectrum Disorders, 27, 44–53. doi:10.1016/j.rasd.2016.04.002

Hojsgaard, S., Halekoh, U., & Yan, J. (2016, September 24). Geepack: Generalized estimating equation package. Retrieved December 19, 2018, from https://CRAN.R-project.org/package=geepack

Hommer, R. E., & Swedo, S. E. (2015, March 1). Schizophrenia and autism—related disorders. Schizophrenia Bulletin, 41 (2), 313–314. doi:10.1093/schbul/sbu188

Hurley, R. S. E., Losh, M., Parlier, M., Reznick, J. S., & Piven, J. (2007). The Broad Autism Phenotype Questionnaire. J Autism Dev Disord, 37 (9), 1679–90.

Hus, V., Gotham, K., & Lord, C. (2014, October). Standardizing ADOS Domain Scores: Separating severity of social affect and restricted and repetitive behaviors. Journal of autism and developmental disorders, 44 (10), 2400–2412. doi:10.1007/s10803-012-1719-1. pmid: 23143131

Hus, V., & Lord, C. (2014, August). The Autism Diagnostic Observation Schedule, Module 4: Revised algorithm and standardized severity scores. Journal of autism and developmental disorders, 44 (8), 1996–2012. doi:10.1007/s10803-014-2080-3. pmid: 24590409

Insel, T. R. (2010, November 10). Rethinking schizophrenia. Nature, 468 (7321), nature09552. doi:10.1038/nature09552

Jiujias, M., Kelley, E., & Hall, L. (2017, December 1). Restricted, repetitive behaviors in autism spectrum disorder and obsessive–compulsive disorder: A comparative review. Child Psychiatry & Human Development, 48 (6), 944–959. doi:10.1007/s10578-017-0717-0

Kanner, L. (1943). Autistic disturbances of affective contact. Nervous Child, 2, 217–50.

Kolvin, I. (1971, April). Studies in the childhood psychoses I: Diagnostic criteria and classification. The British Journal of Psychiatry, 118 (545), 381–384. doi:10.1192/bjp.118.545.381

Kuhn, R., & Cahn, C. H. (2004, September 1). Eugen Bleuler’s concepts of psychopathology. History of Psychiatry, 15 (3), 361–366. doi:10.1177/0957154X04044603

Kushima, I., Aleksic, B., Nakatochi, M., Shimamura, T., Okada, T., Uno, Y., … Ozaki, N. (2018, September 11). Comparative analyses of copy-number variation in autism spectrum disorder and schizophrenia reveal etiological overlap and biological insights. Cell Reports, 24 (11), 2838–2856. WOS:000444267600005. doi:10.1016/j.celrep.2018.08.022

Lieberman, J. A., & First, M. B. (2018, July 19). Psychotic disorders. New England Journal of Medicine, 379 (3), 270–280. doi:10.1056/NEJMra1801490

Lord, C., Elsabbagh, M., Baird, G., & Veenstra-VanderWeele, J. (2018, August 2). Autism spectrum disorder. The Lancet. doi:10.1016/S0140-6736(18)31129-2

Lord, C., Rutter, M., DiLavore, P. C., Risi, S., Gotham, K., & Bishop, S. L. (2012). Autism Diagnostic Observation Schedule Modules 1–4 (2nd). Torrance, CA: Western Psychological Services.

Maijer, K., Begemann, M. J. H., Palmen, S. J. M. C., Leucht, S., & Sommer, I. E. C. (2018, April). Auditory hallucinations across the lifespan: A systematic review and meta-analysis. Psychological Medicine, 48 (6), 879–888. doi:10.1017/S0033291717002367

Marshall, C. R., Howrigan, D. P., Merico, D., Thiruvahindrapuram, B., Wu, W., Greer, D. S., … CNV and Schizophrenia Working Groups of the Psychiatric Genomics Consortium. (2017, January). Contribution of copy number variants to schizophrenia from a genome-wide study of 41,321 subjects. Nature Genetics, 49 (1), 27–35. doi:10.1038/ng.3725

Mazefsky, C. A., Anderson, R., Conner, C. M., & Minshew, N. (2011, March). Child Behavior Checklist scores for school-aged children with autism: Preliminary evidence of patterns suggesting the need for referral. Journal of psychopathology and behavioral assessment, 33 (1), 31–37. doi:10.1007/s10862-010-9198-1. pmid:22661827

McCarthy, S. E., Makarov, V., Kirov, G., Addington, A. M., McClellan, J., Yoon, S., … Sebat, J. (2009). Microduplications of 16p11.2 are associated with schizophrenia. Nature Genetics, 41 (11), 1223–7. doi:10.1038/ng.474

McGlashan, T. H., Miller, T. J., Woods, S. W., Hoffman, R. E., & Davidson, L. (2001). Instrument for the Assessment of Prodromal Symptoms and States. In T. Miller, S. A. Mednick, T. H. McGlashan, J. Libiger, & J. O. Johannessen (Eds.), Early Intervention in Psychotic Disorders (pp. 135–149). NATO Science Series. doi:10.1007/978-94-010-0892-17

McGrath, L. M., Yu, D., Marshall, C., Davis, L. K., Thiruvahindrapuram, B., Li, B., … Scharf, J. M. (2014, August 1). Copy number variation in obsessive-compulsive disorder and tourette syndrome: A cross-disorder study. Journal of the American Academy of Child & Adolescent Psychiatry, 53 (8), 910–919. doi:10.1016/j.jaac.2014.04.022

Meier, S. M., Petersen, L., Pedersen, M. G., Arendt, M. C. B., Nielsen, P. R., Mattheisen, M., … Mortensen, P. B. (2014, November 1). Obsessive-compulsive disorder as a risk factor for schizophrenia: A nationwide study. JAMA Psychiatry, 71 (11), 1215–1221. doi:10.1001/jamapsychiatry.2014.1011

Millan, M. J., Andrieux, A., Bartzokis, G., Cadenhead, K., Dazzan, P., Fusar-Poli, P., … Weinberger, D. (2016, July). Altering the course of schizophrenia: Progress and perspectives. Nat Rev Drug Discov, 15, 485–515. doi:10.1038/nrd.2016.28

Niarchou, M., Chawner, S. J. R. A., Doherty, J. L., Maillard, A. M., Jacquemont, S., Chung, W. K., … van den Bree, M. B. M. (2019, January 16). Psychiatric disorders in children with 16p11.2 deletion and duplication. Translational Psychiatry, 9 (1), 8. doi:10.1038/s41398-018-0339-8

Ooi, Y. P., Rescorla, L., Ang, R. P., Woo, B., & Fung, D. S. S. (2011, September 1). Identification of autism spectrum disorders using the Child Behavior Checklist in singapore. Journal of Autism and Developmental Disorders, 41 (9), 1147–1156. doi:10.1007/s10803-010-1015-x

Owen, M. J., O’Donovan, M. C., Thapar, A., & Craddock, N. (2011, March). Neurodevelopmental hypothesis of schizophrenia. The British Journal of Psychiatry, 198 (3), 173–175. doi:10.1192/bjp.bp.110.084384

R Core Team. (2018). R: A language and environment for statistical computing. Retrieved from https://www.r-project.org

Rapoport, J. L., Giedd, J. N., & Gogtay, N. (2012, December). Neurodevelopmental model of schizophrenia: Update 2012. Molecular Psychiatry, 17 (12), 1228–1238. doi:10.1038/mp.2012.23

Rapoport, J., Chavez, A., Greenstein, D., Addington, A., & Gogtay, N. (2009, January 1). Autism spectrum disorders and childhood-onset schizophrenia: Clinical and biological contributions to a relation revisited. Journal of the American Academy of Child & Adolescent Psychiatry, 48 (1), 10–18. doi:10.1097/CHI.0b013e31818b1c63. pmid: 19218893

Rees, E., Walters, J. T. R., Georgieva, L., Isles, A. R., Chambert, K. D., Richards, A. L., … Kirov, G. (2014, February). Analysis of copy number variations at 15 schizophrenia-associated loci. The British Journal of Psychiatry, 204 (2), 108–114. doi:10.1192/bjp.bp.113.131052. pmid: 24311552

Robinson, D., Hayes, A., Gomez, M., Demeshev, B., Menne, D., Nutter, B., … Werner, K. D. (2018, December 5). Broom: Convert statistical analysis objects into tidy tibbles. Retrieved December 19, 2018, from https://CRAN.R-project.org/package=broom

Rutter, M. (1972 Oct-Dec). Childhood schizophrenia reconsidered. Journal of Autism and Childhood Schizophrenia, 2 (4), 315–337. pmid: 4581613

Rutter, M., Bailey, A., & Lord, C. (2003). Social Communication Questionnaire. Western Psychological Services.

Rutter, M., Le Couteur, A., & Lord, C. (2003). Autism Diagnostic Interview-Revised. Torrance, CA: Western Psychological Services.

Shaffer, D., Fisher, P., Lucas, C. P., Dulcan, M. K., & Schwab-Stone, M. E. (2000, January). NIMH Diagnostic Interview Schedule for Children Version IV (NIMH DISC-IV): Description, differences from previous versions, and reliability of some common diagnoses. Journal of the American Academy of Child & Adolescent Psychiatry, 39 (1), 28–38. doi:10.1097/00004583-200001000-00014

Shank, L. (2011). Mullen Scales of Early Learning. In Encyclopedia of Clinical Neuropsychology (pp. 1669–1671). doi:10.1007/978-0-387-79948-31570

Simons VIP Consortium. (2012, March 22). Simons Variation in Individuals Project (Simons VIP): A genetics-first approach to studying autism spectrum and related neurodevelopmental disorders. Neuron, 73 (6), 1063–1067. doi:10.1016/j.neuron.2012.02.014. pmid: 22445335

Sparrow, S. S., Cicchetti, D. V., & Balla, D. A. (2005). Vineland Adaptive Behavior Scales Vineland-II: Survey Forms Manual. Pearson.

Steinberg, S., de Jong, S., Mattheisen, M., Costas, J., Demontis, D., Jamain, S., … Stefansson, K. (2014, January). Common variant at 16p11.2 conferring risk of psychosis. Molecular Psychiatry, 19, 108–14. doi:10.1038/mp.2012.157

Swets, M., Dekker, J., van Emmerik-van Oortmerssen, K., Smid, G. E., Smit, F., de Haan, L., & Schoevers, R. A. (2014, February 1). The obsessive compulsive spectrum in schizophrenia, a meta-analysis and meta-regression exploring prevalence rates. Schizophrenia Research, 152 (2), 458–468. doi:10.1016/j.schres.2013.10.033

Van Dael, F., van Os, J., de Graaf, R., ten Have, M., Krabbendam, L., & Myin-Germeys, I. (2011, February). Can obsessions drive you mad? Longitudinal evidence that obsessive-compulsive symptoms worsen the outcome of early psychotic experiences. Acta Psychiatrica Scandinavica, 123 (2), 136–146. doi:10.1111/j.1600-0447.2010.01609.x

van Smeden, M., de Groot, J. A. H., Moons, K. G. M., Collins, G. S., Altman, D. G., Eijkemans, M. J. C., & Reitsma, J. B. (2016, November 24). No rationale for 1 variable per 10 events criterion for binary logistic regression analysis. BMC Medical Research Methodology, 16. doi:10.1186/s12874-016-0267-3. pmid: 27881078

Vittinghoff, E., & McCulloch, C. E. (2007, March 15). Relaxing the rule of ten events per variable in logistic and Cox regression. American Journal of Epidemiology, 165 (6), 710–718. doi:10.1093/aje/kwk052

Wechsler, D. (1999). Wechsler Abbreviated Scale of Intelligence. San Antonio, TX: Psychological Corporation.

Weiss, L. A., Shen, Y., Korn, J. M., Arking, D. E., Miller, D. T., Fossdal, R., … Daly, M. J. (2008, February 14). Association between microdeletion and microduplication at 16p11.2 and autism. New England Journal of Medicine, 358 (7), 667–675. doi:10.1056/NEJMoa075974. pmid: 18184952

Wickham, H., & Bache, S. M. (2014, November 22). Magrittr: A forward-pipe operator for R. Retrieved April 9, 2019, from https://CRAN.R-project.org/package=magrittr

Wickham, H., François, R., Henry, L., & Müller, K. (2018, November 10). Dplyr: A grammar of data manipulation. Retrieved December 19, 2018, from https://CRAN.R-project.org/package=dplyr

Wolff, S. (2004, August). The history of autism. Eur Child Adolesc Psychiatry, 13, 201–8. doi:10.1007/s00787-004-0363-5

Zarrei, M., MacDonald, J. R., Merico, D., & Scherer, S. W. (2015, March). A copy number variation map of the human genome. Nature Reviews Genetics, 16 (3), 172–183. doi:10.1038/nrg3871

Zheng, Z., Zheng, P., & Zou, X. (2018, August 1). Association between schizophrenia and autism spectrum disorder: A systematic review and meta-analysis. Autism Research, 11 (8), 1110–1119. doi:10.1002/aur.1977

